# A novel application tunnel in combination with medical training reduces stress induced by frequent intraperitoneal injections and blood sampling in mice

**DOI:** 10.64898/2026.01.08.698459

**Authors:** A. Schlutt, K. Riesner, M. Kochan, A. Meß, D. Jahn, I. Lurje, W. Werner, F. Heymann, Olivia Kershaw, F. Tacke, B. Hiebl, L. Hammerich

## Abstract

Handling-induced stress represents a major burden on laboratory mice and is heavily influenced by the handling technique. As such, tail-handling induces much higher stress than cup- or tunnel-handling, which is further aggravated by interventions, e.g. injections, which require even harsher fixation methods. Previous studies demonstrated that habituation protocols can improve animal welfare during handling and interventions. Here, we developed a medical training program and an alternative fixation method using an application tunnel in the context of a liver cancer model, where frequent intraperitoneal injections over many weeks are needed to induce tumor development. The training regimen consisted of 5 sessions over 2 weeks, which gradually introduced the animals to being touched by handlers and restrained for procedures. The training program was completed once before the start of any interventions, additional training sessions were performed biweekly during the entire course of the experiments. The animals were randomized to receive injections either in the novel application tunnel or using conventional fixation. Training effect was continuously monitored by measuring the latency to interact with the experimenter, the surface body temperature and by movement tracking. The latency to interact rapidly decreased during initial training sessions, and this effect was sustained throughout the course of treatment. Movement tracking demonstrated that mice injected inside the tunnel were more active and returned to their normal behavior after injections faster than conventionally restrained mice. Those mice also showed less signs of defecation and urination. Furthermore, the tunnel had a positive influence on the well-being of mice during blood sampling demonstrated by reduced signs of pain and faster willingness to interact with the handler after the procedure. In conclusion, habituation of the mice to the interventions with medical training and the improved handling procedure during ip injections and blood draws durably reduces stress levels and improves welfare of mice.

## Introduction

Laboratory animals, especially rodents, are still indispensable for medical research to understand the function of genes, the mechanisms of diseases and the effectiveness and toxicities of drugs and chemicals. This is particularly relevant for cancer research, as many hallmarks of cancer can only be fully recapitulated in models using living organisms (1). The house mouse (Mus musculus) is the most frequently used animal species for biomedical research in the European Union; about 1 million mice were used in 2023 for scientific purposes in Germany alone (2) . Directive 2010/63/EU(3) adapted by the European Parliament and the Council of the European Union aims to standardize and harmonize national legal provisions on animal welfare in research. Member States are encouraged to promote research in the field of the 3Rs, which are embedded in article 4 of the Directive (3) . 3R stands for “*Replacement”*, “*Reduction”* and “*Refinement”* as defined by William Russell und Rex Burch’s *„The Principles of Human Experimental Technique*” in 1959 (4). Nowadays, replacement of animal studies with alternative strategies is often viewed as the ultimate goal. However, if replacement by animal-free methods is not possible, refinement methods can be used to improve animal welfare in basic and translational medical research.

Identifying and implementing the most appropriate refinement methods is always dependent on the individual experimental design. It may include enhanced cage enrichment, proven stress-reducing handling methods like cup or tunnel handling or a habituation of the animals to the interventions during experiments in form of a training program. Handling is the procedure most frequently experienced by laboratory animals (during regular husbandry as well as experimental interventions). Especially handling by less-experienced personnel can lead to repeated attempts during handling sessions with an increased risk of being bitten, increased experience of stress and a higher risk of painful mistakes due to wrong restraining or application. Ultimately, handling-induced stress may influence the results of the respective research question(5, 6). Therefore, it appears promising that the implementation of a medical training could be a useful tool to improve animal well-being in a laboratory context since laboratory routine itself is stressful to mice (5). It has been shown that habituation to the method of handling can lower long-term stress and anxiety, as it helps the animals to associate handling with neutral or positive experiences and better tolerate brief unpleasant procedures. (6, 7). Habituation involves repeatedly exposing animals to a non-threatening stimulus such as simple, repetitive handling(8). Currently available protocols usually employ training programs that last 3 to 5 days and sometimes include cognitive enrichment methods such as clicker training as well(6, 9, 10). Handling can also be trained for specific procedures, such as injections or oral gavage(11). It is important to emphasize that the type and frequency of handling depends on the particular experimental setup and any training program must be well thought out to fit the need. Because experimental procedures vary widely, only very limited universal training guidelines exist.

Here, we describe the implementation of a medical training program and a novel application tunnel for injections and blood draws in the context of a mouse model for hepatocellular carcinoma (HCC), which requires repeated intraperitoneal (ip) injections over many weeks. Regarding our training program, the different steps can be adapted depending on the experimental design. In order to determine the training effect, we evaluated several non-invasive stress parameters, including the latency to interact with the experimenter, surface body temperature, corticosterone accumulation in fur as well as the general behavior of the mice. The latency to interact is an easily measurable parameter that can provide immediate information about the stress level and well-being of mice (12), which are flight animals that hide in dangerous situations. The more comfortable a mouse feels, the more social interaction with humans can be observed (13). Another common method to detect acute stress or illness in rodents is measuring the body temperature (14). Since intrarectal measurements can cause additional stress (15), we chose to measure the surface temperature with a contactless infrared thermometer (16). We also measured corticosterone accumulation in fur as an end-point parameter, as it has been shown that corticosterone increases in serum and fur not only due to heightened activity or arousal(17) but also as a response to stress (18) (19), which can be influenced by the handling method (20) (21). Finally, we evaluated the general behavior of the mice after injections by assessing moving patterns and defecation/urination. After blood draws, the mouse grimace scale (MGS) was applied as a readout for pain.

Our data show that habituation of mice to the interventions with medical training and use of the new application tunnel reduced stress levels during ip injections and blood draws as demonstrated by a decreased latency to interact, reduced defecation and urination and lower MGS. In addition, behavioral tracking after ip injections demonstrated that mice injected inside the tunnel were much more active after injections and returned to their normal behavior faster than conventionally restrained mice, even after injection of substances that are known to have a negative effect on well-being themselves. These effects were sustained throughout the course of the experiment. In summary, we established and validated a successful training regimen and a new application method, which are broadly applicable and easy to implement in other studies involving frequent injections and blood draws in mice.

## MATERIALS AND METHODS

### Animals and housing

C57Bl/6J and C57Bl/6N wildtype mice were bred in-house in the experimental area of the Research Institutes for Experimental Medicine (FEM) of Charité - Universitätsmedizin Berlin. Therefore, no acclimatization period to the experimental area was necessary. Mice were kept in groups in type-III-cages at a relatively constant temperature (range 20–24°C) and humidity (range 30–70%). Animals had access to food and water *ad libitum* and cages were changed once per week. All cages were enriched with sawdust bedding, paper nesting material, igloos, opaque tunnels to hide and clear acrylic tunnels, which were also used for contact-free transfer of the mice between cages during the weekly cage changes. Mice were maintained on a 12/12 h light/dark cycle and experiments were conducted between 4 pm and 7 pm when the animals are more active than in the morning hours. All animal experiments were reviewed and approved by the competent local authority (Landesamt für Gesundheit und Soziales Berlin (LAGeSo); approval numbers G 0310/19, G 0112/23, G 0131/23, G 0037/21) and were conducted in accordance with the ARRIVE 2.0 guidelines.

### Hepatocellular carcinoma induction

HCC induction was performed in male mice only, because HCC is much more prevalent in men than women and accordingly, early exposure to the DNA-alkylating agent diethyl nitrosamine (DEN) leads to HCC development in almost 100% of male mice, but only in 30% of female mice. This is thought to be due to estrogen(22, 23). HCC was induced based on established protocols(24–26): male C57Bl/6J mice received a single ip injection of 25mg/kg DEN at the age of 14 days followed by repetitive (twice weekly) injections of carbon tetrachloride (CCl_4_; 0.5ml/kg diluted in corn oil) to induce fibrosis or feeding a Western-style diet (WD) to induce steatosis from the age of 8 weeks (**Figure 1**). All male mice born in our facility received DEN injections and were subsequently randomized to receive CCl_4_ or WD when they were 8 weeks old. The combination of DEN and CCl_4_ resulted in tumor development within 10 weeks (i.e. at 18 weeks of age), the combination of DEN with WD resulted in tumor development within 16 weeks (i.e. at 24 weeks of age). Tumor-bearing mice were again randomized to be treated with an immunotherapy regimen (either a dendritic cell growth factor, fms-like tyrosine kinase 3 ligand (Flt3L), an immune checkpoint inhibitor, or both) administered via ip injection.

**Figure 1.**
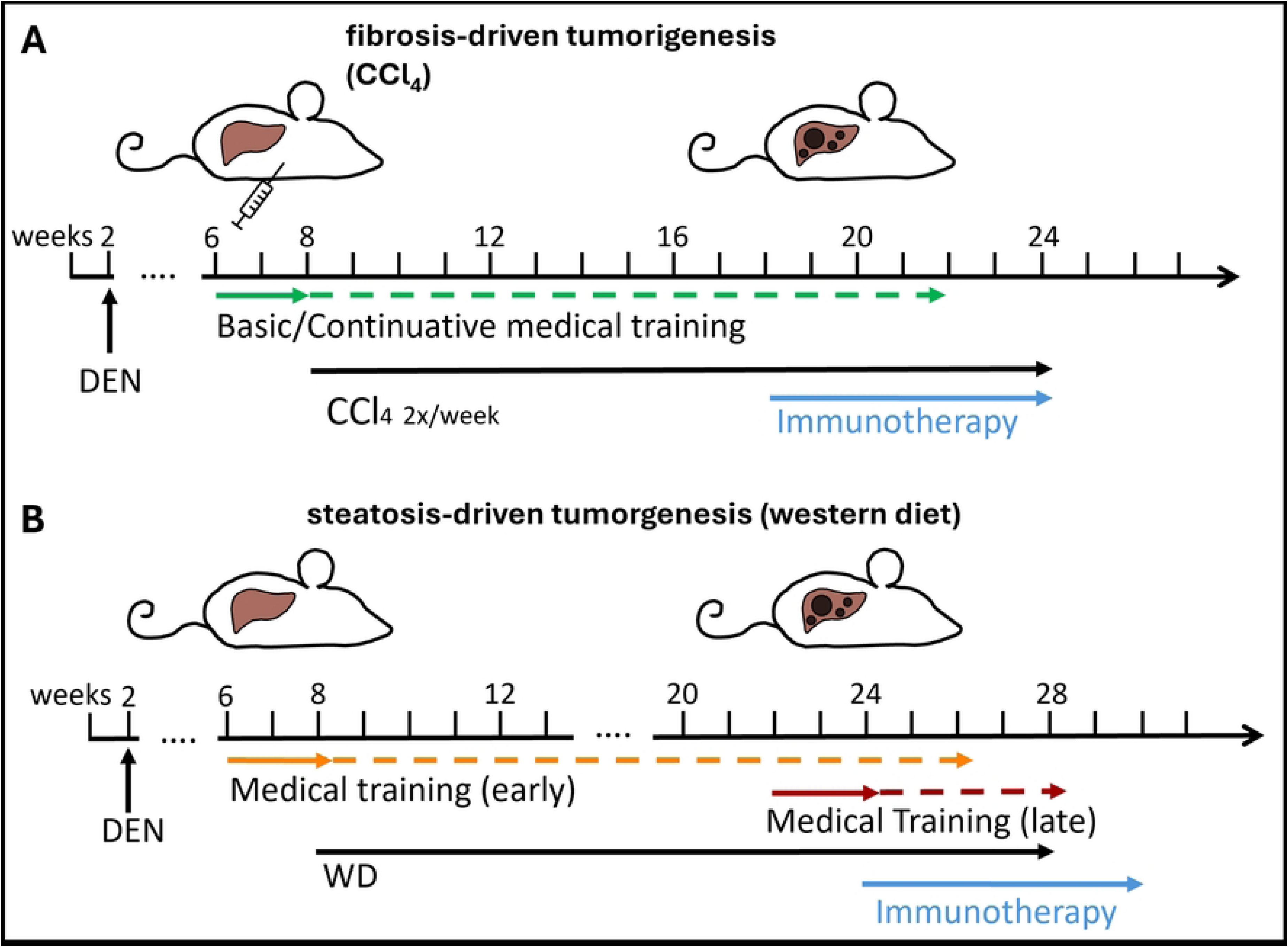
Mouse models. To induce tumorigenesis, mice are injected with diethyl nitrosamine (DEN) at the age of 14 days and then subjected to carbon tetrachloride (CCl_4_) injections to induce fibrosis or western diet (WD) feeding to induce steatosis. (A) fibrosis model: Basic medical training program starts at 6 weeks of age, CCl_4_ injections start at 8 weeks of age. Training is continued (2 sessions every other week) until immunotherapy treatments commence at 18 weeks after tumors have developed. (B) steatotis model: Basic medical training program starts either at 6 weeks of age („early“; 2 weeks before WD feeding starts at 8 weeks) and is continued (2 sessions every other week) until immunotherapy treatments commence at 24 weeks or starts at 22 weeks („late“; 2 weeks before immunotherapy treatments).

### Experimental groups

Following the 3R principle of reduction, this welfare study was conducted exclusively on animals already undergoing HCC induction in other ongoing projects. All animals undergoing HCC induction during the course of this study were included. Within the CCl_4_ model, mice were randomized to receive medical training and/or ip injections in the novel application tunnel (**Table 1**, groups 2-5). Within the WD model, mice were randomized to receive medical training either early (at 6 weeks, ie. 2 weeks before beginning of WD feeding) or late (at 20 weeks, ie. 2 weeks before ip injections begin) (**Table 1**, groups 6-8). As a baseline control group for biochemical measurements, 22-weeks old healthy mice without any interventions (no disease induction) were included as well (**Table 1**, group 1). Because animals entered the experiments at 2 weeks (=before weaning), animal numbers differed between cages depending on the number of male pups per litter. In general, mice in the same experimental groups were housed together. Therefore, we treated cage as the experimental unit, unless otherwise stated. Blinding of the experimenters during training and sample/data collection was not possible, because all animals had to be trained repeatedly and many measurements were taken during or immediately after training. This required the experimenters to be aware of the group allocation of the animals at any given time. However, experimenters were blinded to sample allocation during end-point sample analysis whenever possible (eg. corticosterone measurements, pathological examination).

**Table 1.** Overview of experimental groups.

| # | Disease model | Treatment | Training regimen | Injection method | Cage | Mouse | Corticosterone | Tracking | Pathology |
| --- | --- | --- | --- | --- | --- | --- | --- | --- | --- |
| 1 | Untreated<br>(n=6) | None | None | None | C1 | M1 | x |  |  |
|  |  |  |  |  | C2 | M2 | x |  |  |
|  |  |  |  |  |  | M3 | x |  |  |
|  |  |  |  |  |  | M4 | x |  |  |
|  |  |  |  |  | C3 | M5 | x |  |  |
|  |  |  |  |  |  | M6 | x |  |  |
| 2 | CCl<sub>4</sub><br>(n=15) | Ip-injections twice a week starting at week 8 (CCl<sub>4</sub>) and daily at week 18 (Immunotherapy) | None | Conventional | C4 | M7 | x |  |  |
|  |  |  |  |  |  | M8 | x |  |  |
|  |  |  |  |  | C5 | M9 | x |  |  |
|  |  |  |  |  |  | M10 |  |  |  |
|  |  |  |  |  |  | M11 | x |  |  |
|  |  |  |  |  | C6 | M12 | x | x |  |
|  |  |  |  |  |  | M13 | x | x |  |
|  |  |  |  |  |  | M14 | x | x |  |
|  |  |  |  |  |  | M15 | x | x |  |
|  |  |  |  |  |  | M16 | x | x |  |
|  |  |  |  |  | C7 | M17 |  | x |  |
|  |  |  |  |  |  | M18 |  | x |  |
|  |  |  |  |  |  | M19 |  | x |  |
|  |  |  |  |  |  | M20 |  | x |  |
|  |  |  |  |  |  | M21 |  | x |  |
| 3 | CCl<sub>4</sub>+T<br>(n=10) | Ip-injections twice a week starting at week 8 (CCl<sub>4</sub>) and daily at week 18 (Immunotherapy) | None | Tunnel | C8 | M22 | x |  | x |
|  |  |  |  |  |  | M23 | x |  |  |
|  |  |  |  |  |  | M24 | x |  |  |
|  |  |  |  |  |  | M25 |  |  | x |
|  |  |  |  |  | C9 | M26 | x |  |  |
|  |  |  |  |  |  | M27 | x |  |  |
|  |  |  |  |  |  | M28 |  |  |  |
|  |  |  |  |  | C10 | M29 |  |  | x |
|  |  |  |  |  |  | M30 |  |  | x |
|  |  |  |  |  |  | M31 |  |  | x |
| 4 | CCl<sub>4</sub>+MT<br>(n = 23) | Ip-injections twice a week starting at | Basic training from week 6, | Conventional | C11 | M32 |  |  |  |
|  |  |  |  |  |  | M33 |  |  |  |
|  |  | week 8 (CCl<sub>4</sub>) and daily at week 18 (Immunotherapy) | followed by continuous training (2 sessions every other week) |  | C12 | M34 | x |  |  |
|  |  |  |  |  |  | M35 | x |  |  |
|  |  |  |  |  |  | M36 | x |  |  |
|  |  |  |  |  |  | M37 | x |  |  |
|  |  |  |  |  |  | M38 | x |  |  |
|  |  |  |  |  | C13 | M39 |  |  |  |
|  |  |  |  |  |  | M40 |  |  |  |
|  |  |  |  |  |  | M41 | x |  |  |
|  |  |  |  |  | C14 | M42 |  |  |  |
|  |  |  |  |  |  | M43 |  |  |  |
|  |  |  |  |  |  | M44 |  |  |  |
|  |  |  |  |  |  | M45 |  |  |  |
|  |  |  |  |  |  | M46 |  |  |  |
|  |  |  |  |  | C15 | M47 | x |  |  |
|  |  |  |  |  |  | M48 | x |  |  |
|  |  |  |  |  |  | M49 | x |  |  |
|  |  |  |  |  |  | M50 | x |  |  |
|  |  |  |  |  | C16 | M51 |  |  |  |
|  |  |  |  |  |  | M52 |  |  |  |
|  |  |  |  |  |  | M53 |  |  |  |
|  |  |  |  |  |  | M54 |  |  |  |
| 5 | CCl<sub>4</sub>+MT+T (n=14) | Ip-injections twice a week starting at week 8 (CCl<sub>4</sub>) and daily at week 18 (Immunotherapy) | Basic training from week 6, followed by continuous training (2 sessions every other week) | Tunnel | C17 | M55 | x |  |  |
|  |  |  |  |  |  | M56 | x |  |  |
|  |  |  |  |  |  | M57 | x |  |  |
|  |  |  |  |  | C18 | M58 | x | x |  |
|  |  |  |  |  |  | M59 | x | x |  |
|  |  |  |  |  |  | M60 | x | x |  |
|  |  |  |  |  |  | M61 | x | x |  |
|  |  |  |  |  |  | M62 | x | x |  |
|  |  |  |  |  | C19 | M63 |  |  | x |
|  |  |  |  |  |  | M64 |  |  | x |
|  |  |  |  |  | C20 | M65 | x | x |  |
|  |  |  |  |  |  | M66 | x | x |  |
|  |  |  |  |  |  | M67 |  | x |  |
|  |  |  |  |  |  | M68 |  | x |  |
| 6 | WD<br>(n=9) | WD from week 8,<br>daily ip-injections<br>at week 24<br>(Immunotherapy) | None | Conventional | C21 | M69 | x |  |  |
|  |  |  |  |  |  | M70 | x |  |  |
|  |  |  |  |  |  | M71 | x |  |  |
|  |  |  |  |  | C22 | M72 | x |  |  |
|  |  |  |  |  |  | M73 | x |  |  |
|  |  |  |  |  |  | M74 | x |  |  |
|  |  |  |  |  |  | M75 | x |  |  |
|  |  |  |  |  |  | M76 | x |  |  |
|  |  |  |  |  |  | M77 | x |  |  |
| 7 | WD+MT,<br>early<br>(n=29) | WD from week 8,<br>daily ip-injections<br>at week 24<br>(Immunotherapy) | Basic training<br>from week 6,<br>followed by<br>continuous<br>training (2<br>sessions every<br>other week) | Conventional | C23 | M78 |  |  |  |
|  |  |  |  |  |  | M79 |  |  |  |
|  |  |  |  |  |  | M80 |  |  |  |
|  |  |  |  |  |  | M81 |  |  |  |
|  |  |  |  |  |  | M82 |  |  |  |
|  |  |  |  |  |  | M83 |  |  |  |
|  |  |  |  |  | C24 | M84 |  |  |  |
|  |  |  |  |  |  | M85 |  |  |  |
|  |  |  |  |  |  | M86 |  |  |  |
|  |  |  |  |  |  | M87 |  |  |  |
|  |  |  |  |  | C25 | M88 | x |  |  |
|  |  |  |  |  |  | M89 | x |  |  |
|  |  |  |  |  |  | M90 | x |  |  |
|  |  |  |  |  |  | M91 | x |  |  |
|  |  |  |  |  | C26 | M92 | x |  |  |
|  |  |  |  |  |  | M93 |  |  |  |
|  |  |  |  |  |  | M94 |  |  |  |
|  |  |  |  |  |  | M95 |  |  |  |
|  |  |  |  |  |  | M96 | x |  |  |
|  |  |  |  |  | C27 | M97 |  |  |  |
|  |  |  |  |  |  | M98 | x |  |  |
|  |  |  |  |  |  | M99 |  |  |  |
|  |  |  |  |  |  | M100 |  |  |  |
|  |  |  |  |  |  | M101 | x |  |  |
|  |  |  |  |  | C28 | M102 | x |  |  |
|  |  |  |  |  |  | M103 |  |  |  |
|  |  |  |  |  |  | M104 |  |  |  |
|  |  |  |  |  |  | M105 |  |  |  |
|  |  |  |  |  |  | M106 | x |  |  |
| 8 | WD+MT<br>late<br>(n= 21) | WD from week 8,<br>daily ip-injections<br>at week 24<br>(Immunotherapy) | Basic training<br>from week 22 (2<br>weeks before ip-injections begin) | Conventional | C29 | M107 | x |  |  |
|  |  |  |  |  |  | M108 | x |  |  |
|  |  |  |  |  |  | M109 | x |  |  |
|  |  |  |  |  |  | M110 | x |  |  |
|  |  |  |  |  |  | M111 | x |  |  |
|  |  |  |  |  | C30 | M112 | x |  |  |
|  |  |  |  |  |  | M113 | x |  |  |
|  |  |  |  |  |  | M114 | x |  |  |
|  |  |  |  |  |  | M115 | x |  |  |
|  |  |  |  |  | C31 | M116 |  |  |  |
|  |  |  |  |  |  | M117 |  |  |  |
|  |  |  |  |  |  | M118 |  |  |  |
|  |  |  |  |  | C32 | M119 |  |  |  |
|  |  |  |  |  |  | M120 |  |  |  |
|  |  |  |  |  |  | M121 |  |  |  |
|  |  |  |  |  | C33 | M122 |  |  |  |
|  |  |  |  |  |  | M123 |  |  |  |
|  |  |  |  |  |  | M124 |  |  |  |
|  |  |  |  |  |  | M125 |  |  |  |
|  |  |  |  |  |  | M126 | x |  |  |
|  |  |  |  |  |  | M127 |  |  |  |

### Medical training program

All animals undergoing HCC induction were included in the medical training program and analyzed according to their group assignment (no exclusions of animals occurred). The training program included a basic training consisting of 5 training sessions within 2 weeks (**Table 2**, **Figure 2**) and was started at the age of 6 weeks, ie. 2 weeks before the beginning of CCl_4_ injections or WD feeding to induce liver cancer (**Figure 1**). The amount of training sessions was chosen based on previous studies demonstrating training success after completing 5 training sessions with mice (27). At the end of every training session mice were given treats (Kellogg’s fruit loops®) in the home cage and training sessions were 2 to 3 days apart to allow time for recovery. The same Vetbed was used for all mice in one cage (but not shared between cages) and reused over several training days. Vetbeds were cleaned after contamination with urine/feces. A detailed description of the training sessions included considerations for handler behavior, criteria of success and how to respond to animal behavior can be found in Table 2. In general, handlers performing the training should stay calm, avoid fast movement and not put any pressure on the animals (eg. corner them) as this can lead to negative associations.

**Figure 2.**
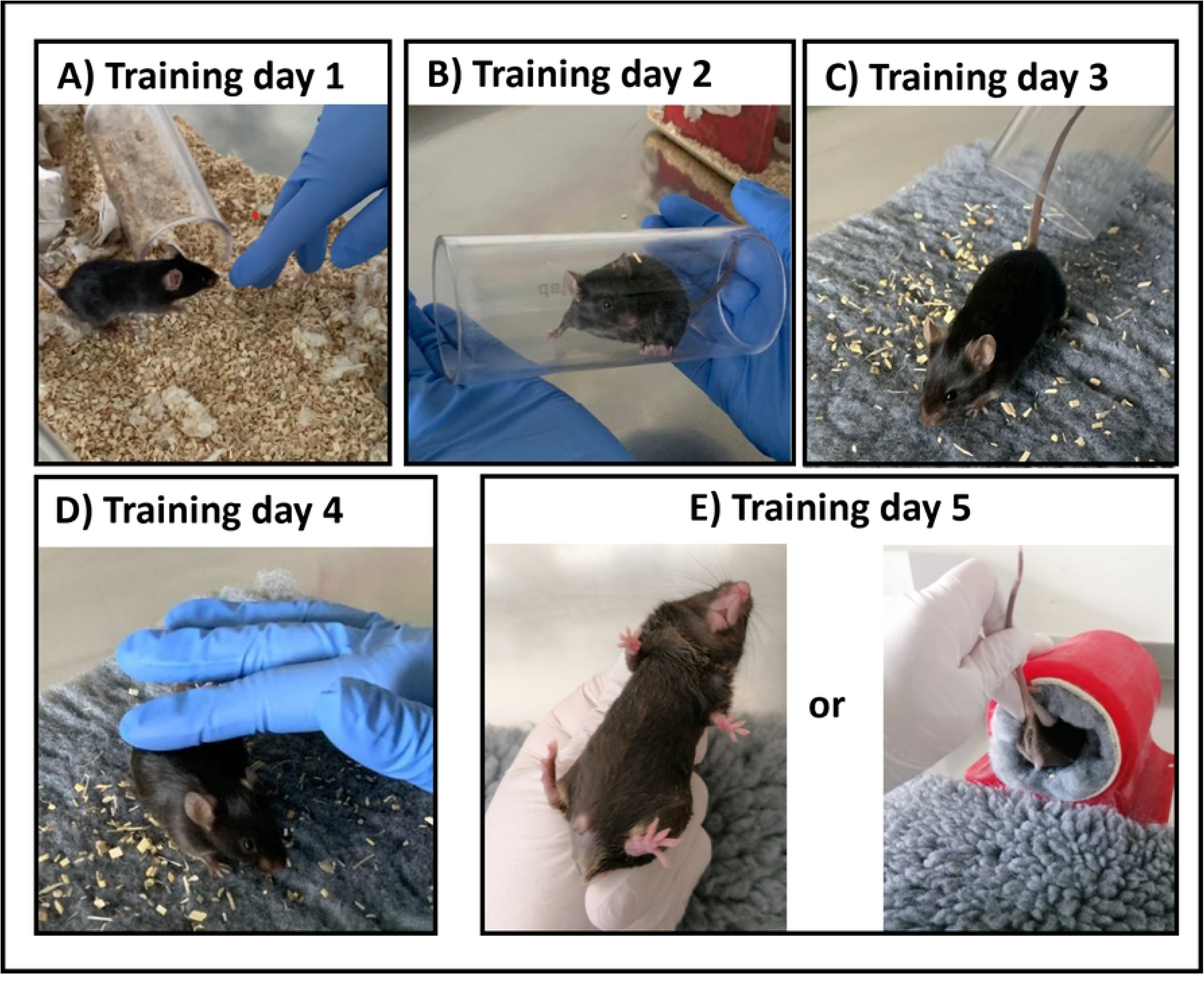
Medical Training Program. (A) Day 1: Gloved hand is reached inside the cage. (B) Day 2: mice are directed into acrylic handling tunnels and held for about 30 seconds before being released back into the cage. (C) Day 3: mice are transfered onto a Vetbed outside of the cage using the acrylic handling tunnel. (D) Day 4: same as day 3, in addition mouse is gently touched for about 30 seconds. (E) Day 5: fixation method for injections is performed once without injection.

**Table 2.** Overview of the basic training program. 5 sessions are spread over 2 weeks with 2-3 days between each session. In general, signs of stress or anxiety, like the latency to interact or defecation/urination, should decrease over the training sessions and stay low.

| Training day | Training program | Important considerations for handler behavior | Criteria of success | response to animal behavior |
| --- | --- | --- | --- | --- |
| 1<br>(Fig. 2a) | Handler opens the cage, reaches the gloved hand inside and waits for voluntary interaction. If possible, handler tries to carefully touch the mouse. | • Reach hand in one corner rather than the middle of the cage to not pressure the animals.<br>• Hold hand still and wait for voluntary interaction | Animals approach hand voluntarily | If no voluntary interaction occurs within 5 minutes, session should be terminated and repeated on the next day. |
| 2<br>(Fig. 2b) | Handler opens the cage and directs the mice one by one into handling tunnels, which will be lifted. The handler closes both ends of the tunnel with the hands to prevent animals from walking back out and holds handling tunnel in the air for 30 seconds, before placing the mice | • Guiding the mice into the tunnel can take longer for some mice in the beginning, e.g. 30 - 60 s.<br>• When guiding the mice into the tunnel, it's important to not "chase" the mice with the tunnel and | Mice enter tunnel without force. | • If mice refuse to enter the tunnel, terminate session after 2min.<br>• In the beginning, mice often show signs of anxiety in the tunnel, such as defecation and urination. This is not a reason to terminate the training session but this type of behavior should |
|  | back into their home cages. | to not trap them in a corner to force them to enter the tunnel. This would lead to negative associative sensitization. |  | decrease with training progression. |
| **3**<br>(Fig. 2c) | Similar procedure as on day two, but the handling tunnels are used to transfer the mice onto a Vetbed outside of the cage. After being transferred, the handler tries to touch the animals gently for about 30 seconds, before transferring them back into the cage using the handling tunnel. | • Aim to let the animals get in contact with the handler first (e.g. sniffing) before touching.<br>• Because touching the back area can cause more sensitive reactions, it is suggested to start with the head area of the animals. | • Animals leave tunnel voluntarily and explore Vetbed<br>• Animals tolerate brief touching | • The mice should leave the tunnel voluntarily. If not, the handler can slightly increase the angle from the tunnel to the vet bed so that the mouse slides out very gently. In our experience, this works with less stress if the mouse is allowed to slide out caudal end first. |
| **4**<br>(Fig. 2d) | Same as on day 3, with a focus on touching the mice in different areas (eg. neck, back, tail). | • Aim to let the animals approach the handler first before touching. | • Animals show less avoidant behavior (eg. flight or freezing) | • If animals act nervous and try to escape, it can be helpful to offer them the tunnel, so they may hide for a moment |
|  |  |  | when being touched<br>Animals show less signs of defecation or urination | before being let out onto the VetBed again |
| 5<br>(Fig. 2e) | Training of the handling for intraperitoneal injection: either conventional scruffing or directing the mice into the injection tunnel and holding the tail/sacrum for a few seconds. No actual injection is performed. | Use a syringe without needle to touch the mice in the lower caudal area where injection would normally be performed<br>Do not restrain the animals for too long (about the time that would be needed for performing an injection) | Mice enter injection tunnel voluntarily/can be guided in easily | Mice may show signs of anxiety such as defecation and urination (especially when being scruffed). This should decrease with training progression.<br>If mice cannot be guided into the injection tunnel easily, terminate session after 5min and repeat on the following day. |

**Table 3.** Overview of experimental groups to evaluate effect of restraining method on stress levels after blood sampling.

| # | Disease model | Procedure | Training regimen | Restraining Method |
| --- | --- | --- | --- | --- |
| 1 | Untreated (n=8) | Blood draw from tail vein | None | Conventional Restrainer |
| 2 | Untreated (n=7) | Blood draw from tail vein | None | Tunnel |

### Conventional Handling (scruffing) for injections

Mice were immobilized on a grid by pushing down the lower abdomen and extremities with one hand while pulling back (scruffing) the fur and skin in the neck area between thumb and forefinger of the other hand. Mice were turned around to expose the abdomen and the tail fixated between the palm and the little finger of the same hand holding on to the neck (**Figure 2E**), leaving the other hand free for injections (28).

### Application tunnel for intraperitoneal injections

The application tunnel was 3D-printed in house, details on the design and manufacturing can be found in supplementary methods. The size of the tunnel was dependent on the age of mice; for mice under 10 weeks a tunnel with an inner diameter of 7 cm, for older mice of 8 cm was used. The length of the tunnel was 10 cm. A rolled-up piece of a washable and autoclavable Vetbed, which the mice were introduced to during the medical training, was placed inside the tunnel. Since studies have shown that mice feel safer on groundings which have similar color to their fur(29), a grey Vetbed was used (**Figure 3A**). The same Vetbed was used for all mice housed together in one cage, but changed between cages. After each day of injection both the Vetbed and tunnel were cleaned.

**Figure 3.**
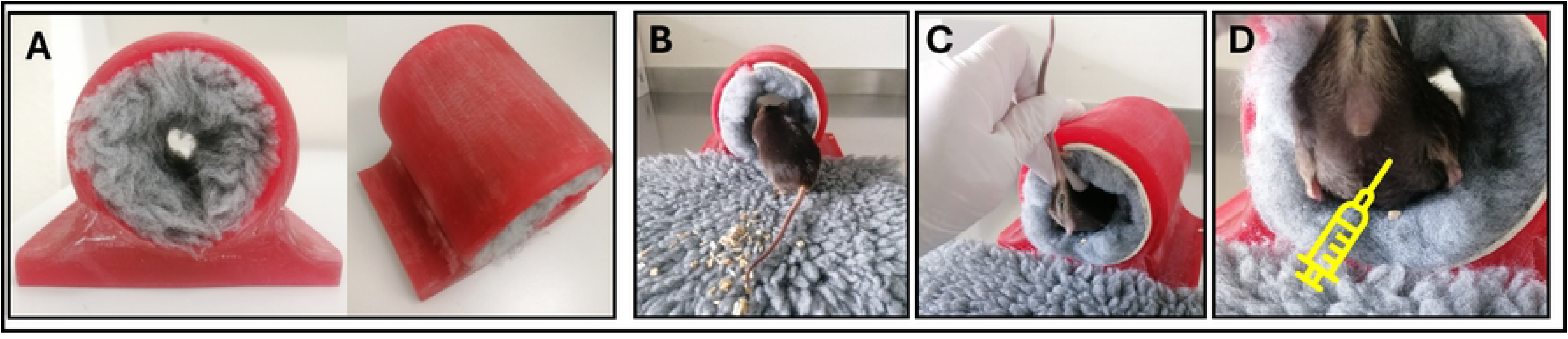
Injection Tunnel. (A) 3D-printed tunnel with Vetbed. (B) Mouse entering tunnel. (C) Soft pressure is applied on the sacral area and tail fixed between forefinger and thumb. (D) View on caudal abdominal region for ip injection and location of injection (syringe)

### Injections inside the tunnel

Figure 3 shows the procedure of ip injection using the tunnel (a link to a video demonstrating the injection procedure can be found in supplementary methods). Mice were removed from their home cages using the acrylic tunnel already known from the medical training and placed on a Vetbed outside of the cage. In our experience, the mice often enter the application tunnel voluntarily and instinctively as it offers a place to hide. If not, mice were gently directed to enter the injection tunnel (Figure 3B) The handler carefully held the tail’s base between forefinger and thumb to prevent that the mouse is walking too deep inside. By applying soft pressure on the sacral area with the middle finger of the same hand, the mouse is immobilized (Figure 3C). This fixation allows a good view on the caudal abdominal region where ip injection is performed (Figure 3D). After completing the injection, the injection tunnel was placed inside the home cage to allow mice to return to their home cage without additional handling.

### Latency to interact with the experimenter

Social interaction or lack thereof can be used as a parameter for stress (27). For the purpose of this study, we defined the latency to interactas the amount of time required for a mouse to voluntarily get into contact with the handler after completing a training session. To measure this, mice were placed back into their home cage after each training session or after blood draws and one gloved hand of the handler was reached inside the cage but did not move. A timer was used to measure the time it took for the animal to voluntarily interact with the gloved hand again (eg. sniff). Each mouse was assessed individually, but with the co-housed animals present in the home cage. Cage means were treated as the experimental unit, unless stated otherwise.

### Body temperature

Surface body temperature was determined non-invasively using an infrared thermometer (GT-101, Geratherm Medical AG, Geratal, Germany) by holding the thermometer over the back region of the mice (2-3mm away). Temperature measurements were performed immediately at the end of each training session and again 15 minutes after completing the training session and transferring the mice back into their home cages. Each mouse was assessed individually, but with the co-housed animals present in the home cage. Cage means were treated as the experimental unit.

### Subcutaneous tracking chips and movement tracking

For movement tracking, a Radio Frequency Identification (RFID) Chip (ID162-PM) was implanted subcutaneously in the caudal abdominal region left of the Linea alba, using the IntelliCage® Injektor (Cat.No. D-290000-IC-50705). The procedure was performed under short-term isoflurane anesthesia. The 2 x 2 mm wound channel was closed using a single staple with a 5/0 Monocryl suture. Cages were placed on Trafficage devices (TSE Systems, Berlin, Germany) to monitor movement of the animals inside the cages. Every cage was divided into 6 areas and each time an animal moved from one area to another was recorded as an event (**Fig. S2**). Measurements were analyzed using Microsoft Access. Co-housed mice were in the same experimental groups and tracked at the same time; cage means were treated as the experimental unit.

### Blood sampling from the tail vein

Healthy untrained female C57Bl/6N mice (not undergoing HCC induction) were used to assess stress levels after blood draw from the tail vein. Mice were randomized to undergo blood sampling either in a conventional restrainer or in the injection tunnel. In both cases the lateral tail vein was scored with a needle and one drop of blood collected into a microtube. Stress parameters were assessed immediately after releasing mice from the restrainer or the tunnel: pictures to assess MGS were taken on the Vetbed as soon as the mice exited the restrainer or tunnel; latency to interact was measured directly after placing the animals back into their home cages individually. Therefore, Individual mice were treated as the experimental units for stress parameters taken after blood sampling. Two animals had to be excluded from the study, because blood draw was not successful on the first try and animals had to undergo restraining procedure twice, which could potentially falsify the results due to higher stress levels.

### Mouse grimace scale

The MGS was first introduced in 2010 to detect facial pain expression and categorization in mice (30). Since then, it is a widely used parameter in pain and stress assessment (31). For classification of animal welfare the MGS can be used as a scoring system for the characterization of reaction to pain by analyzing expression of facial muscles (31, 32) and is a useful tool to determine the stress level in mice, eg. after repeated anesthesia (33). The scores were determined using photographs (n= 2-3 per mouse) taken directly after the procedures before placing the animals back into their home cages. Therefore, individual animals were treated as the experimental unit in this case. Two observers with an experienced background on mice behavior scored three different facial expression parameters: Orbital tightening, nose bulge, and ear position. We did not assess whisker position and cheek bulge as this was challenging due to the dark fur color of our mice (a limitation already noticed in other animal welfare studies (34, 35)). The parameters were scored on a three-point scale from 0 to 2 (0 = not present, 1 = moderately present, 2 = obviously present). The three facial scores were then added up and averaged (mean MGS scores). Each animal was scored by both observers independently and the 2 MGS scores averaged to reach one final MGS score per animal.

### Pathological examination

3 conventionally injected mice and 4 mice injected in the tunnel were randomly selected to be sampled for histopathological examination of the injection site. At sacrifice, tissue samples of about 2 x 1 cm including peritoneum, abdominal wall and skin were collected from the caudal abdominal region where mice had received injections and stored in 4 % paraformaldehyde solution until further processing. Samples were paraffin embedded, cut into 5 µm sections and stained with hematoxylin and eosin at the Institute of Veterinary Pathology, Freie Universität Berlin. Histopathological examination was performed by an EBVS® European Specialist in Veterinary Pathology.

### Fur collection and Corticosterone analysis

Fur samples were collected by carefully cutting the hair on the back of the mice with fine scissors as close as possible to the skin. Samples were wrapped in aluminum foil and stored at room temperature in the dark to avoid light exposure. Steroid extraction and quantification was performed by *Dresden Lab Service GmbH* (Dresden, Germany) as previously described (36, 37). Briefly, the fur samples were washed by shaking them in 2.5 mL isopropanol for 3 min at room temperature. After drying for 12 hours, 10 mg of non-pulverized hair was transferred to a 2 ml centrifuge tube. After adding 50 μl of internal standard and 1,8 mL Methanol, the sample was incubated for 18 h. Thereafter, the sample was centrifuged at 10000 rpm for 2 min and 1 ml of the supernatant was transferred to a new 2 ml tube. Alcohol was evaporated at 65°C and under a constant stream of nitrogen. The dried samples were resuspended in water and analyzed via liquid chromatography–tandem mass spectrometry (LC-MS/MS) assay with atmospheric pressure chemical ionization (APCI) positive mode on a Shimadzu LC-20AD HPLC unit with a Shimadzu SIL-20AC autosampler and a Shimadzu CTO-20AC column temperature oven.

### Statistical Analysis

Statistical analysis was performed using GraphPad Prism 10. Normality of data sets was tested using Shapiro-Wilk test. Comparison of two groups was performed using paired or unpaired Student’s T test for normally distributed data and Mann Whitney test for non-parametric data. Comparison of more than two groups was performed using repeated measures one-way ANOVA with Geisser-Greenhouse correction followed by Dunnett’s multiple comparison test (comparing every mean to a control mean) or repeated measures 2-way ANOVA followed by Šidák’s multiple comparison test, as indicated. P-values under 0.05 were considered significant. Two mice had to be excluded from experiments due to technical issues during blood draw (see above), but no additional data points or outliers were removed during analysis.

### Data availability

This study protocol has been preregistered on animalstudyregistry.org and will be made publicly available upon publication. Technical diagrams and 3D-printing files for manufacturing the tunnel as well as a video demonstrating the injection procedure have been deposited on the Open Science Framework platform (OFS.io) and will be made publicly available upon publication under this link (view-only for peer review): https://osf.io/vsjnb/?view_only=ff970c7b95104b0888e26b00f43cc30c The data that support the findings of this study are not openly available due to reasons of sensitivity and are available from the corresponding author upon reasonable request. Data are located in controlled access data storage at Charité Universitätsmedizin Berlin.

## RESULTS

### Latency to interact

As a first and very immediate parameter to determine training effect, we measured the latency to interactafter every training session. As expected, the latency to interactwas quite high and very variable in untrained mice not used to the experimenter, but decreased in all groups already after the first day of training (Figure 4 **A+B**). This effect was sustained even after the beginning of frequent ip injections (between training day 5 and 6 for CCl_4_ mice and between training day 17 and 18 for WD mice). Furthermore, in the CCl_4_ model we did not observe any differences between mice injected using the conventional scruffing method or the injection tunnel (Figure 4A).

**Figure 4.**
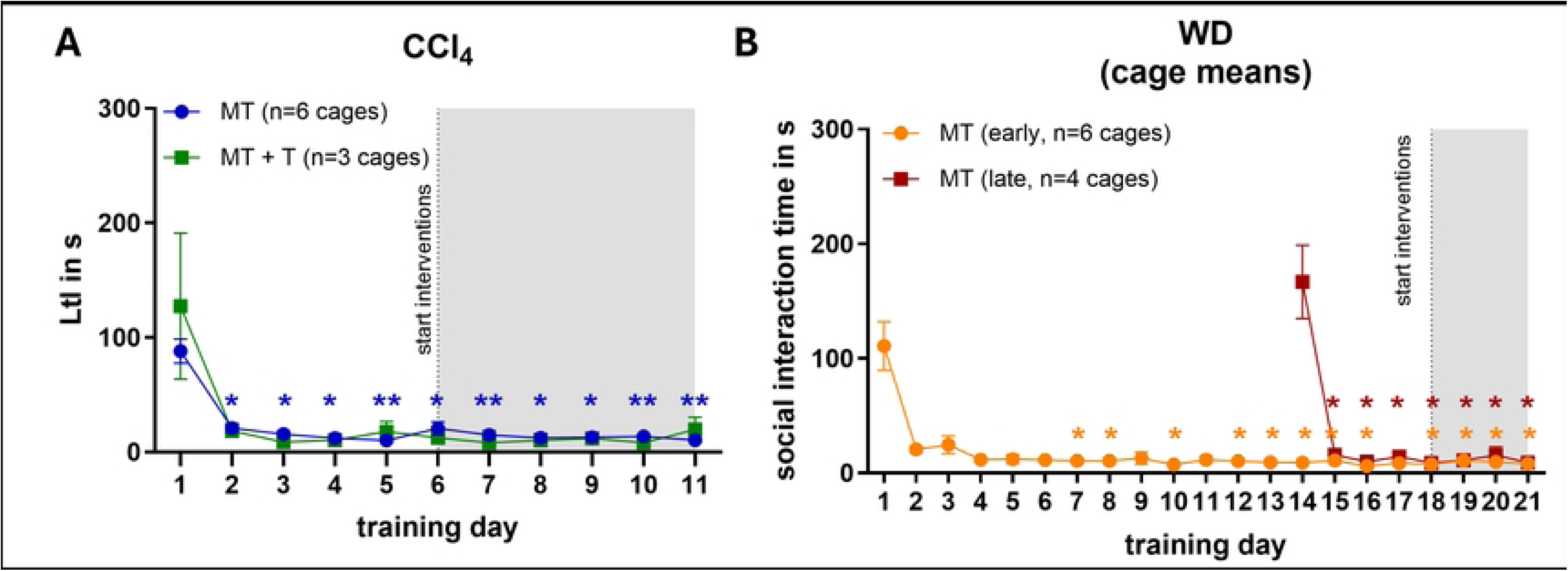
Latency to interact. Mice were subjected to the medical training regimen described in Fig.1 starting at the age of 6 weeks. After each training session the Latency to interact (LtI) was recorded for every animal individually, but with co-housed animals present in the home cage. Cage means (average LtI of all mice in one cage) were treated as the experimental unit. (A) Latency to interact of CCl_4_ mice undergoing medical training only (MT) or medical training with tunnel injection (MT+T), (B) Latency to interact of WD mice receiving early (MT early) vs. late (MT late) medical training. Data are displayed as mean +/- SEM. Repeated Measures One-way ANOVA with Geisser-Greenhouse correction and Dunnett’s multiple comparisons test (all time-points compared to first day of training). . *=p<0.05, **=p<0.01, ***=p<0.001, ****=p<0.0001

In the WD model, we also compared mice that received an early basic training starting at the age of 6 weeks followed by a continuous training (MT early) with mice that received a late training starting at the age of 22 weeks (WD+ MT late) (Figure 4B). As described above, we observed a strong decrease in the latency to interactafter the first training day in both groups, which did not increase again after the beginning of the ip injections, indicating that the age of the mice did not have an influence on the training effect.

### Body temperature

Next, we assessed the surface body temperature immediately and 15min after each training session. We expected to observe a higher temperature in stressed mice due to the stress-induced hyperthermia response(38, 39). Hence, body temperature should be higher immediately after the training sessions as compared to 15min after returning to their home cage and successful stress reduction via medical training should result in a lower surface body temperature over time. However, we observed the opposite in the CCl_4_ + MT group (Figure 5A). The temperature right after the training session was lower (mean minimum 35,33°C, maximum 35,61°C) compared to the temperature 15 minutes after (mean minimum 35,66°C, maximum 35,82°C). Of note, this difference was could be observed at every timepoint during the course of the experiment, but did not reach significance in most cases. No remarkable changes were detectable depending on training success or age. Similar results were obtained for MT + T mice with minimal surface measurements of 34,8°C and maximum measurements of 35,69°C right after the training and a minimum of 35,35°C and a maximum of 35,86°C after 15 minutes recovery (Figure 5B). It is important to note that CCl_4_ -treated mice were handled inside a biosafety cabinet with a ventilation system with high air flow due to the hazardous properties of CCl_4_, which results in a rather cold surrounding atmosphere.

**Figure 5.**
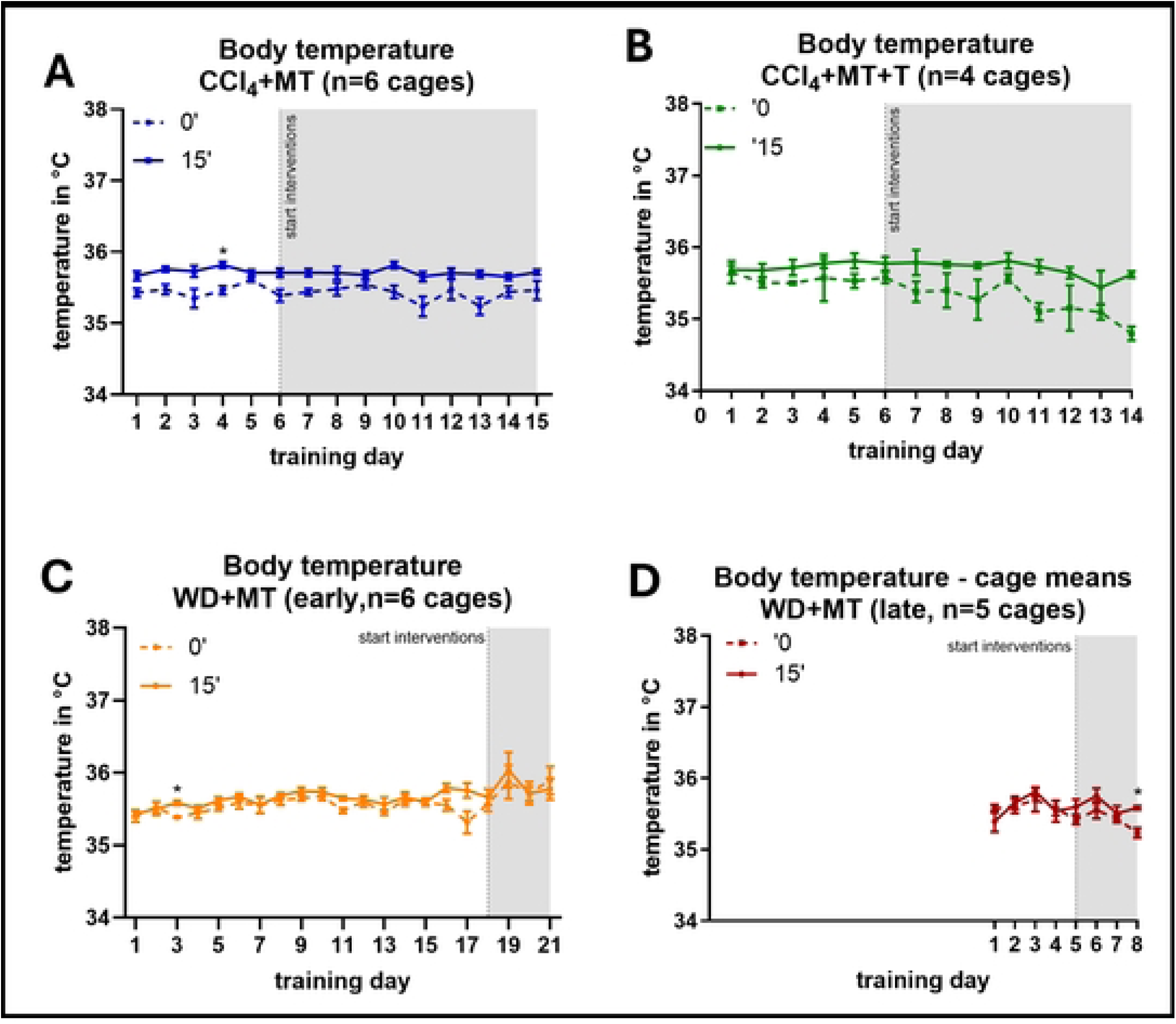
Body temperature. Body temperature measured immediately (0’) and 15 minutest (15’) after training sessions. Body temperature was measured for every animal individually, but with co-housed animals present in the home cage. Cage means (average body temperature of all mice in one cage) were treated as the experimental unit. (A) CCl4 mice with medical training (CCl4 +MT), (B) CCl4 mice with medical training and tunnel injection, (C) WD mice with early medical training (WD+MT early) and (D) WD mice with late medical training (WD+MT late). Data are displayed as mean +/- SEM. 2-way ANOVA with Šidák’s post-test. *=p<0.05

In comparison, mice of the WD-model (Figure 5C**+D**) showed no differences of surface body temperature right after (Mean WD early = 35,61 °C and WD late = 35,43°C) and 15 minutes after the medical training session (Mean WD early = 35,7 °C and WD late = 35,44°C). Those mice underwent medical training under the same conditions as their housing conditions (20-24°C). The animals had an increased body temperature over their training sessions and lifetime, and the WD early group displayed a temporary rise in temperature right after immunotherapy onset.

### Behavioral assessment after ip injections

In previous studies, we noticed that mice often show an apathic behavior directly after ip injections, especially after the injection of CCl_4_. CCl_4_ intoxication is known to cause symptoms like headaches, nausea and abdominal pain in humans (40), which may well explain the side-effects we observe in our mice: about 5 to 10 min after the injection, the animals start showing apathic behavior including closed eyes, curved backs, laid-back ears and above all, severely restricted movement for 30 to 60 minutes. Since we observed that this behavior seemed less pronounced when the mice were injected inside the novel injection tunnel, we assessed the activity of the mice after CCl_4_ ip injections using subcutaneous RFID chips. For each mouse, the number of movements was collected over a period of three hours following ip injection either in the tunnel or using the conventional scruffing method for several (up to 19) time points per mouse (Figure 6). To generate a baseline, the same mice were tracked after cage cleaning and medical training sessions (**Fig. S3A**). Since mice in the same experimental groups were housed together and assessed at the same time, we compared the cage means at each time point rather than the movements of individual mice. In general, all mice were less active after ip injections than after training sessions, but mice injected inside the tunnel were more active after the injections than conventionally injected mice (Figure 6A). In addition, tunnel injected mice seemed to recover faster as well (Figure 6B). During the first hour after injection, the count of movements did not differ between both groups, however, during the second and third hour the tunnel injected group showed significantly more movement than conventionally injected mice. Both groups had a decreased movement again into the third hour after injection, possibly due to the handler having left them alone for quite some time by then, but tunnel injected mice were still more active than the conventionally injected group. Importantly, the positive effect on the activity of the mice did not seem dependent on the injected substance, as we observed the same trend after ip injection of our immunotherapy drug (**Flt3L, Fig**. S**3B-C**).

**Figure 6.**
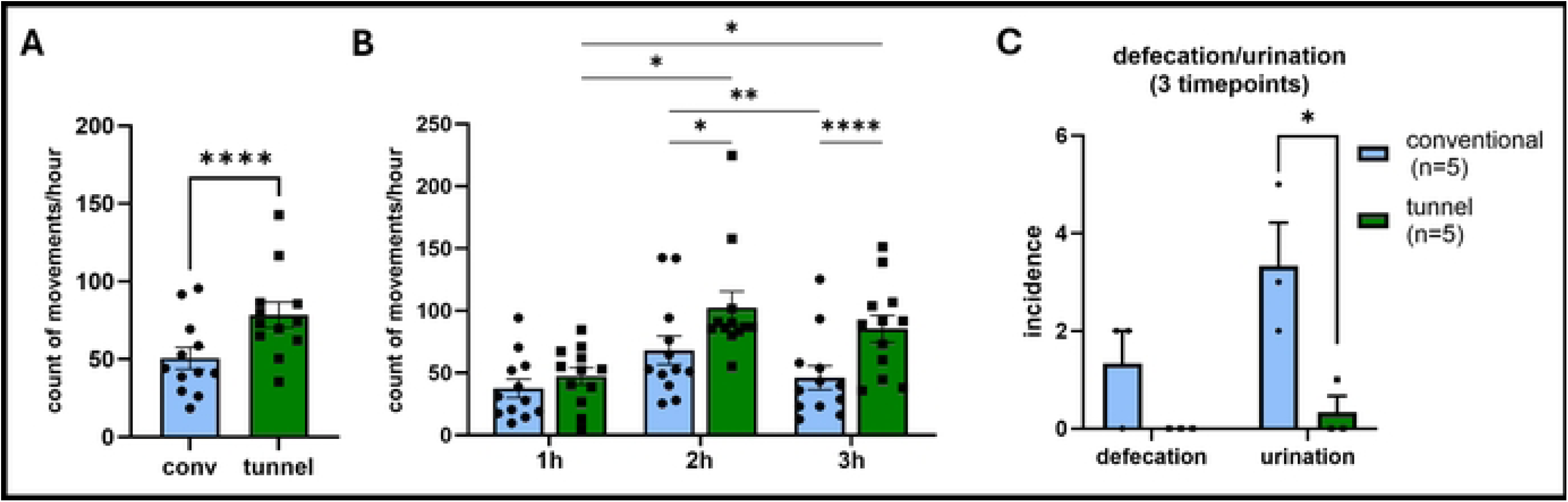
Behavioral tracking after ip injections. Mice were implanted a subcutaneous RFID chip in the abdominal region. After ip injections using either conventional scruffing or the novel injection tunnel, cages were placed on a traffic cage system and movements of the mice were recorded for a total of 3 hours after each injection. Because co-housed mice were tracked together, cage means (average count of movements of all mice in one cage per timepoint) were treated as the experimental until. Conventional: n=10 mice in 2 cages; tunnel: n=9 mice in 2 cages. (A) Count of movements after tunnel or conventional injection of CCl_4_ measured over 3h at 12 different timepoints during the course of the experiment. Every data point represents the average count of movement per hour of all mice in the group at one experimental timepoint. Paired Student’s t test; (B) Count of movements after tunnel or conventional injection stratified by the first three hours after CCl_4_. Repeated measures two-way ANOVA with Geisser-Greenhouse correction and Šidák’s post-test; (C) Incidence of urination and defecation during injection (each data point represents the sum of events in each group at 3 different experimental time points). Unpaired Student’s t test. Data are displayed as mean +/- SEM. *=p<0.05, **=p<0.01, ***=p<0.001, ****=p<0.0001

### Defecation and Urination

We also recorded cases of urination and defecation during the injection procedure in two groups of mice, which received CCl_4_ injections either conventionally or in the injection tunnel. During the first 3 injection timepoints (ie. the first 3 injections after completing the training regimen), we detected 10 cases of urination and 4 cases of defecation from conventionally injected mice (n=5) compared to no defecation and only one case of urination from mice injected inside the tunnel (Figure 6C).

### Pathological examination

To ensure that the altered positioning of the animals during injection did not result in injuries to the application area we also examined the injection sites of 3 randomly selected conventionally and 4 randomly selected tunnel-injected mice histopathologically (**Fig. S4**). All mice showed moderate to severe inflammation of the skeletal muscles and, in some cases, seroma-like cystic cavities, which are likely the result of the regular and frequent injection needle puncture. However, in the pathological examination, no differences were observed between the two injection groups in a blinded assessment, suggesting that injection inside the tunnel did not cause any additional injuries.

### Blood sampling

The encouraging results of the movement tracking and behavior of the mice during and after ip injections prompted us to also test the injection tunnel for blood sampling from the tail vein using needle puncture. The existing restrainers to immobilize the mice during this procedure often cause signs of fear like vocalization, defecation and urination. Therefore, we assessed the stress level of mice after blood draw inside the restrainer in comparison to mice after blood draw using the injection tunnel. For this purpose, we used healthy untrained mice in order to assess the effect of the tunnel without any confounders from the disease induction or the training. The latency to interact measured right after blood draw was significantly lower for the tunnel-handled mice (mean = 13,14 s) compared to restrainer-handled mice (mean = 41,5 s) (Figure 7A). We also noticed variation in facial expression and general habitus. Pictures taken right after the blood draw (**Fig. S5**) were assessed for orbital tightening, nose bulge and ear position on a categorical scale from 0 to 2. Tunnel-handled mice displayed a significantly lower MGS score compared to restrained mice (Figure 7B). Taken together, these results indicate that restraining mice in the tunnel also increases welfare during and after blood sampling from the tail vein.

**Figure 7.**
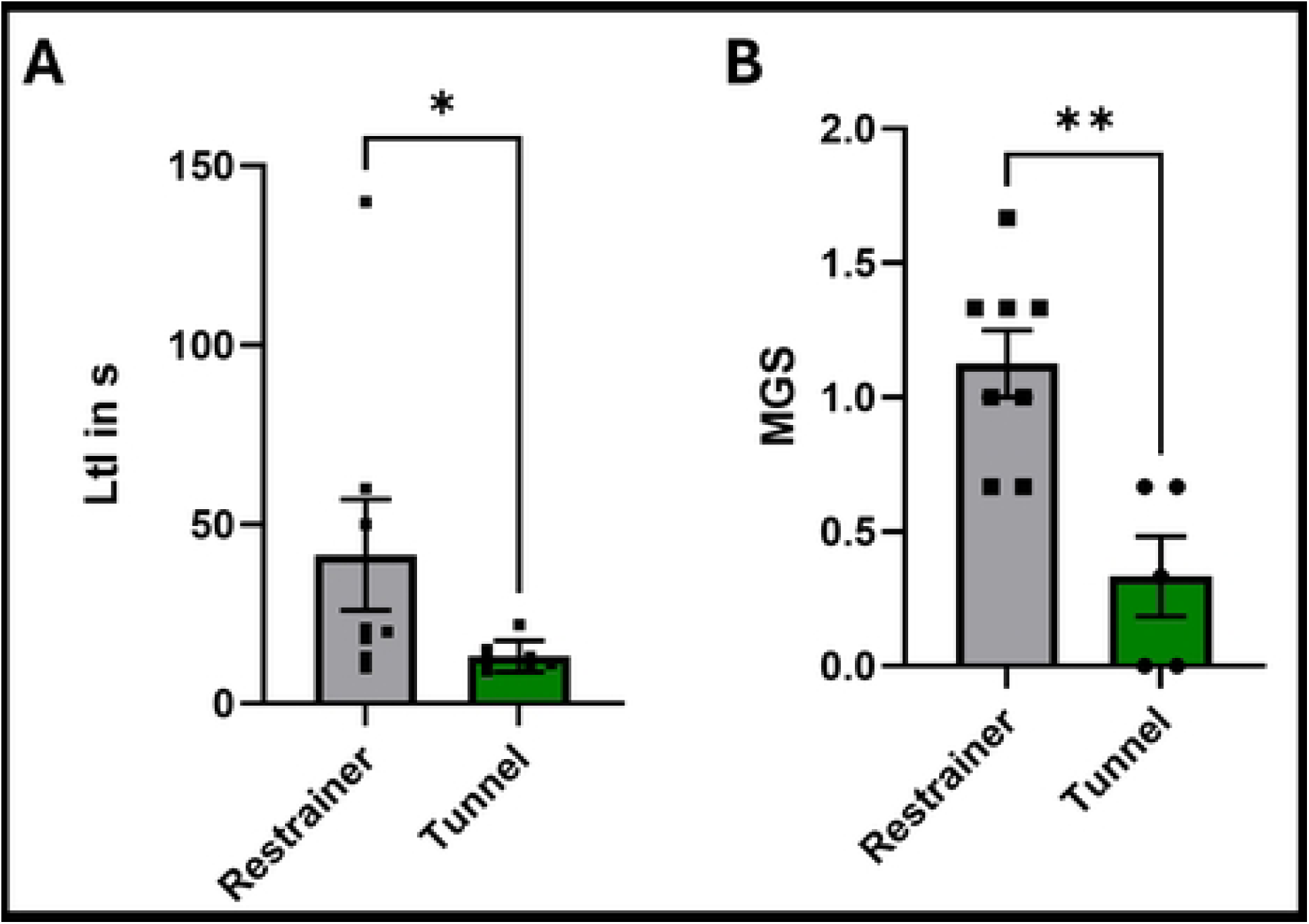
Social interaction time and grimace scale after blood draw from the tail vein. Mice were subjected to blood sampling either inside a conventional restrainer (n=8) or the novel injection tunnel (n=7). (A) Latency to interact (LtI) measured immediately after blood draw. Every animal was assessed individually with no other animals present in the home cage. Mann Whitney test. (B) Facial expression assessed by mouse grimace scale (MGS) immediately after blood draw. Mice were assessed individually before being returned to the home cage. Unpaired Student’s t test, Data are displayed as mean +/- SEM. *=p<0.05, **=p<0.01

### Corticosterone accumulation in fur

As a long-term stress parameter, we assessed corticosterone accumulation in the fur at the age of 22 weeks (CCl_4_ model) and 26 weeks (WD model) and compared corticosterone levels to 22-weeks old healthy untrained mice without any disease induction or interventions. We were not able to detect any significant differences in either model (**Fig. S6A+B**), with a tendency towards lower corticosterone levels in WD mice in the early training group, which also received a continuous training throughout the whole experiment (for about 20 weeks). This may indicate that the medical training has a positive influence on the welfare of the mice even during regular housing without interventions. This effect was not observed in the late training group.

## Discussion

Our study shows that habituation of mice to interventions with medical training in combination with the use of a novel application tunnel reduces stress levels during frequent ip injections and blood draws. Similar to what other studies have shown (9, 27, 41), a training frequency of 5 training days was sufficient for us to get the animals used to the handling procedure, e.g. for ip injections. Neither the timepoint of the training start within the experiment nor the beginning of the injections had an influence on the training success. As expected, measuring the latency to interact with the experimenter as an immediate stress parameter was very helpful to determine the effect of the training. Since this parameter decreased rapidly after only one training session and remained constant after that, it is possible that this change in approach behavior was supported by familiarity with the test from day 2 onwards, which would suggest that less training still maintains the same benefits. As each training session creates some level of stress on the animals, achieving the desired effect with as few training sessions as possible is highly desirable and would also save considerable lab resources (experimenter time). However, we noticed that it is essential for the animals to be introduced to the injection tunnel during the training program in order to facilitate them entering the tunnel without force. Therefore, the sessions for training of the injection procedure should not be omitted. Otherwise, the time saved on training would be offset by the additional time needed to convince animals to enter the tunnel and the added stress of using force might negate the purpose of the training. In contrast to the latency to interact, the measurement of body temperature showed no changes over the course of the medical training sessions and rather seemed dependent on external influences as working under ventilation systems (resulting in lower surrounding temperature) seemed to distort the values strongly. The risk of temperature loss for mice with a relatively large surface area compared to their body weight is known (42). Therefore, the reduced surface temperature observed in CCl_4_ -treated animals was likely due to the surrounding atmosphere in the biosafety cabinet rather than stress-related. In addition, the increase in body temperature observed in WD animals over time was likely due to the diet-induced obesity and decreasing body surface to weight ratio (43) and the temperature peak in the WD early group right after immunotherapy seemed to be more influenced by immunotherapy itself than training effort (44). Therefore, it might have been more informative to test both CCl_4_ and WD mice in similar environmental conditions, however, this was not possible due to animal facility regulations (due to its hazardous properties, CCl_4_ -treated mice had to be kept in a separate room and cages could only be opened under a biosafety cabinet to protect experimenters). Nevertheless, since we were still able to detect rather small temperature deviations (between 0 and 15 min after training in cold environments), surface measurements can generally be a good alternative to intrarectal measurements, which has been suggested by other studies as well (16, 45). It is important to keep in mind that there is a big variety in surface temperature depending on the location of measurement (46), which can falsify the results due to vasoconstriction in the periphery as a reaction to stress and pain. Measuring the temperature in proximal areas and with a tissue contact probe (thermometer measures directly on the surface), as we performed here, leads to more accurate and consistent results compared to a contactless measurement (47).

In mice, handling procedures during laboratory routine itself, which includes lifting the mice up, cleaning and moving the cages, can already trigger fear and pain (5). Invasive interventions, like injections, which are associated with harsh restraining are maximizing those effects. It has been shown that ip injections with saline only can increase stress parameters like fecal corticosterone metabolites (46). Catching mice by the tail can cause pain and fear and many studies demonstrated that tail handling should be exchanged by tunnel or cup handling (13, 27) as even frequent interventions like ip injections couldn’t minimize the positive effects of giving up tail handling (12, 48). In line with this, we observed that not only the method of catching and transferring mice, but also the type of restraining during injection appears to have a major impact. The measurement of movement patterns after ip injections showed that mice that experience less restraining injection had a significantly faster recovery period. Importantly, this effect seemed to be independent of the injected substance. Furthermore, we noticed that use of the injection tunnel can also reduce stress in researchers, especially those inexperienced in handling and fixation of animals. Conventional restraining methods can lead to stress caused by fear of getting bitten while scruffing the animal, which is aggravated by failed attempts and repeated tries to achieve correct positioning of the mouse. Increased stress levels of the researchers transfer to the animals and heightens their fear, which makes restraining even more difficult. The animals included in this study were all handled by experienced handlers to exclude level of experience as a confounder, but many researchers, including some that were newly trained, reported decreased hesitation in approaching the animals and performing injections.

We also observed reduced signs of pain after blood sampling from the tail vein inside the new handling tunnel, which we assessed by taking the MGS immediately after the procedure. Mice handled in a conventional restrainer showed higher MGS compared to those, which experienced the same procedure less restrained in our injection tunnel. This is in accordance with a previous study that demonstrated pain differences based on facial expression during ear-tagging depending on the handling method (tunnel or tail). Tunnel handled mice showed significantly lower MGS after ear-tagging (49), suggesting that handling method plays an important role in the response of mice to stressful and painful procedures. Still, the experience of the scorers can lead to significant variability in MGS results (31) and needs to be taken into account. Already existing approaches in half-automated evaluation of MGS are promising tools for standardized assessment of facial expressions (50) and could allow the use by non- or less-experienced experimenters.

In our study, we could not measure any significant changes in the end-point stress parameter corticosterone in the fur of our mice. Neither medical training nor tunnel injection were able to lower corticosterone levels of the mice in the CCl_4_ groups and only a tendency could be observed in the WD groups. It is unclear, whether this means that chronic stress levels in our mice were unchanged or whether corticosterone levels were influenced by any other experiment-related conditions. It is known that corticosterone levels show higher variability in grouped mice, potentially caused by hierarchy with the dominant animal in the cage showing the highest levels (21). The frequent handling of the CCl_4_ mice can lead to increased aggressiveness and dominance behavior, as the animals constantly have to fight out and defend their hierarchy when reassembling/relocating in their cages, which may cause higher variability of corticosterone levels in individual mice. Additionally, it should be noted that sampling the fur can be falsified by various circumstances. Individual results can be falsified by grooming each other in groups and transferring the corticosterone-containing saliva of the mice to each other. This problem could be circumvented by measuring corticosterone metabolites in urine or feces, but these need to be collected using special metabolic cages, which can lead to increased acute stress levels due to single-housing and absence of nesting material and housing and therefore falsify results (51). In addition, corticosterone is underlying a circadian rhythm with a peak in the evening hours during the awakening response of the nocturnal animals, which further complicates standardized measurements (17). Our results are in line with a recent study in rats that challenged the general concept of hair corticosterone as a representative of chronic stress and concluded that glucocorticoid levels in fur can only indicate recent stress as they diffuse out of hairs again within a few days after a stress stimulus(52). This might explain why our end-point measurements were not able to show any differences between the groups. Further studies with more frequent sampling during the course of experiments would be needed to confirm this.

In conclusion, our study shows that the medical training program is able to habituate the mice to human contact and reduce handling-associated stress during interventions. In addition, the use of the new application tunnel seems to significantly mitigate the negative influence of procedures like injections and blood sampling and further reduced signs of stress, fear and pain. Therefore, our medical training protocol provides a good concept to increase and monitor animal welfare in the context of long-term experiments with frequent ip injections, which can easily be adapted depending on the experimental design. The application tunnel is easily customizable to individual needs as well. It can be printed on basic 3D-printer models using low-cost UV-cured resin making it accessible to a wide scientific community even in limited resource settings. Given the large number of rodents used in biomedical research worldwide, our injection tunnel has the potential to make a significant and lasting impact on laboratory animal welfare.

## ACKNOWLEDGMENTS

We thank our animal welfare officer Dr. Juliane Unger for her support in designing and implementing this study and all members of the Charité 3R Refinement Taskforce for helpful discussion.

## SUPPORTING INFORMATION CAPTIONS

Figure S1. Technical diagram of the application tunnel

Figure S2. (A) Traffic cage system; (B) detection plates with 6 areas to detect movements

**Figure S3. Behavioral tracking after ip injections or cage cleaning.** Mice were implanted a subcutaneous RFID chip in the abdominal region and cages placed on a traffic cage system to record movements of the mice for a total of 3 hours. Every data point represents the average count of movement of all mice in the group at one experimental timepoint. (A) Count of movements after cage cleaning or medical training sessions at 4 different timepoints during the course of the experiment; (B) Count of movements after tunnel (n=4 mice) or conventional (n=5 mice) injection of immunotherapy measured over 3h at 8 different timepoints during the course of the experiment. Paired Student’s t test; (C) Count of movements after tunnel or conventional injection stratified by the first three hours after ip injection. Data are presented as mean +/- SEM. *=p<0.05

Figure S4. Representative H&E pictures of abdominal wall after injection inside the application tunnel (n=4) or using conventional scruffing (n=3)

Figure S5. Pictures used for mouse grimace scale assessment after blood draw in conventional restrainer (A) or tunnel (B)

Figure S6. Corticosterone accumulation in fur.

(A) Untreated, n= 6 mice in n=3 cages; CCl_4_, n= 10 mice in n=2 cages; CCl_4_ + MT, n = 10 mice in n=3 cages; CCl_4_ + MT+ T, n = 10 mice in n=3 cages; CCl_4_ + T, n = 5 mice in n=2 cages;

(B) Untreated, n=6 mice in n=3 cages; WD, n=9 mice in n=2 cages; WD+MT(early), n=10 mice in n=4 cages; WD+MT(late), n=10 mice in n=3 cages.

CCl_4_=carbon tetrachloride; WD=western diet

